# The consequences of tetraploidy on *Caenorhabditis elegans* physiology and sensitivity to chemotherapeutics

**DOI:** 10.1101/2023.06.06.543785

**Authors:** Kelly R. Misare, Elizabeth A. Ampolini, Hyland C. Gonzalez, Kaitlan A. Sullivan, Xin Li, Camille Miller, Bintou Sosseh, Jaclyn B. Dunne, Christina Voelkel-Johnson, Kacy L. Gordon, Jessica H. Hartman

**Author notes:** Equal contribution (Kelly R. Misare, Elizabeth A. Ampolini).

## Abstract

Polyploid cells contain more than two copies of each chromosome. Polyploidy has important roles in development, evolution, and tissue regeneration/repair, and can arise as a programmed polyploidization event or be triggered by stress. Cancer cells are often polyploid. *C. elegans* nematodes are typically diploid, but stressors such as heat shock and starvation can trigger the production of tetraploid offspring. In this study, we utilized a recently published protocol to generate stable tetraploid strains of *C. elegans* and compared their physiological traits and sensitivity to two DNA-damaging chemotherapeutic drugs, cisplatin and doxorubicin. As prior studies have shown, tetraploid worms are approximately 30% longer, shorter-lived, and have a smaller brood size than diploids. We investigated the reproductive defect further, determining that tetraploid worms have a shorter overall germline length, a higher rate of germ cell apoptosis, more aneuploidy in oocytes and offspring, and larger oocytes and embryos. We also found that tetraploid worms are modestly protected from growth delay from the chemotherapeutics but are similarly or more sensitive to reproductive toxicity. Transcriptomic analysis revealed differentially expressed pathways that may contribute to sensitivity to stress. Overall, this study reveals the phenotypic consequences of whole-animal tetraploidy in *C. elegans*.

## INTRODUCTION

Polyploidy—the presence of more than two full sets of chromosomes in a cell—can be brought about during stress to the cell/tissue/organism or arise due to programmed developmental events. Programmed polyploidization has been widely recognized as important in evolution and fitness (Van de Peer et al., 2017), development (Anatskaya and Vinogradov, 2022), tissue repair (Bailey et al., 2021), and regeneration (Øvrebø and Edgar, 2018). In vertebrates, most tissues are mainly diploid, but a high frequency of polyploidy occurs in vascular smooth muscle (McCrann et al., 2008), brain (Jungas et al., 2020), heart (Derks and Bergmann, 2020), and liver (Sladky et al., 2021, Zhang et al., 2019).

In both heart and liver, polyploidy occurs in the forms of mononucleated and multinucleated cells and has suggestive ties to regenerative potential of these tissues. Interestingly, there is wide species-level variation in the extent of heart polyploidy. Mammals exhibit 90-99% polyploidy in the adult heart while zebrafish and newts have just 1-2% polyploid cardiomyocytes. It has been suggested that high rates of polyploidy in the heart may be responsible for the lack of regeneration in this tissue in mammals (Derks and Bergmann, 2020). By contrast, the liver contains approximately 30-50% polyploid hepatocytes in adult humans and 90% polyploid hepatocytes in adult mice (Zhang et al., 2019), and an impressive capacity for regeneration has been observed in both species.

In addition to developmentally programmed polyploidy, spontaneous polyploidization can occur as a result of stress or injury. Liver regeneration after a partial hepatectomy of two-thirds of the liver results in increased polyploidy through mechanisms distinct from those acting during development (Miyaoka et al., 2012). Other stressors can drive polyploidization in liver, including non-alcoholic fatty liver disease (Gentric et al., 2015) and hepatotoxicant exposures such as carbon tetrachloride (Zhang et al., 2019). There is evidence that ploidy changes in the liver are linked to hepatocellular carcinogenesis (Wang et al., 2021) and can drive recurrence/relapse of some tumors (White-Gilbertson and Voelkel-Johnson, 2020). For example, Polyploid Giant Cancer Cells (PGCC) are a heterogenous, senescent population of cells that contain a >4n genome, evade radiation or chemotherapeutic treatments, and can later repopulate the tumor and metastasize (White-Gilbertson and Voelkel-Johnson, 2020). PGCCs are thought to form due to stress from hypoxia, interstitial fluid pressure in tumors, or cancer chemo- and radiation therapies themselves. The development of PGCCs supports the notion that polyploid cells have a higher capacity to respond to and survive in stressful environments. Collectively, diverse studies of polyploidization suggest that it may be a conserved mechanism for stress mitigation.

Studies dating back over 70 years establish that full-body polyploidization can be induced in *Caenorhabditis elegans* by stressing the animals via starvation or heat shock (Nigon, 1949). Recently, a new method for inducing and culturing stable polyploid *C. elegans* lines was developed (Clarke et al., 2018). The method uses RNAi to knock-down a gene encoding a meiosis-specific cohesion protein, *rec-8*, for two generations and produces tetraploid progeny in the third generation. Tetraploid worms are longer than diploid worms, which allows for visual selection of tetraploids and the establishment of stable lines with 100% transmission of tetraploidy. *C. elegans* is a particularly attractive model to study the cellular and organismal effects of polyploidy given the genetic tools available, its fully sequenced and highly annotated genome, and the ability to grow large, genetically homogenous populations under different experimental conditions. Furthermore, the transparent body of *C. elegans* allows visualization of tissue architecture and fluorescent stains and proteins in living animals.

In this study, we use this powerful polyploid *C. elegans* model to further our understanding of the physiological effects of polyploidy. We first evaluated the physiological and metabolic consequences of tetraploidy in worms by measuring their body size, lifespan, transcriptional profiles, and mitochondrial respiration. We then explored the relative sensitivity of these tetraploid worms (compared to diploid) when exposed to genotoxic chemotherapeutics cisplatin and doxorubicin. We tested two exposure paradigms: a developmental exposure and its effect on growth beginning at the L1 larval stage, and a late L4 larval exposure and its effect on reproduction. Together, the results of our study reveal key differences between diploid and tetraploid *C. elegans* and lay the foundation for future work in the study of polyploidy—especially interactions of ploidy and resilience to genotoxic stress—using *C. elegans* as a model.

## MATERIALS AND METHODS

### *C. elegans* Strains

The N2 (wild-type, diploid) *C. elegans* strain was provided by the Caenorhabditis Genetics Center (CGC), which is funded by NIH Office of Research Infrastructure Programs (P40 OD010440). Stable tetraploid N2 *C. elegans* lines were generated using methods previously published (Clarke et al., 2018). In short, wild-type N2 worms were fed HT115 *E. coli* transformed to contain *rec-8* RNAi to knock down the gene. *rec-8* is required for proper pairing and disjunction of chromosomes during meiosis and inhibition results in *C. elegans* polyploidy within two generations. Tetraploid worms in the third generation were identified by visual inspection due to their *lon* phenotype (they are 20-30% longer than diploid worms). Single tetraploid worms were allowed to self-fertilize and the degree of tetraploidy was determined in the offspring. Tetraploidy was confirmed by counting DAPI stained chromosomes in unfertilized oocytes of gravid adult worms using a Leica DMI-8 Thunder Imager inverted microscope. Once stable tetraploid lines were established with 100% tetraploid offspring, all worms were maintained on OP50 K-agar plates (Boyd et al., 2012) at 15°C.

### Chemotherapeutics and Dosing *C. elegans*

The chemotherapeutics (Cisplatin, Teva Pharmaceuticals and Doxorubicin, Hikma Pharmaceuticals) used in this study were obtained from the Medical University of South Carolina Hospital Pharmacy at concentrations of 1 mg/mL for cisplatin and 2 mg/mL for doxorubicin. *C. elegans* were exposed to external doses of 500, 250, 100, 50, 25, and 0 μM cisplatin and 250, 100, 50, 25, 5, and 0 μM doxorubicin diluted in 9 mg/mL sodium chloride solution pH 3.6 (the diluent in the commercial infusions). Briefly, 1.2 mL of the cisplatin solutions and 1 mL of doxorubicin solutions were added to 6 cm OP50 K-agar plates and allowed to absorb/dry at 33°C for approximately 1 hour, or until the surface was completely dry. Appropriate stage *C. elegans* were then cultured on these plates for 24-48h (depending on the experiment).

### Size and Growth Assessments

Diploid and tetraploid *C. elegans* populations were synchronized via egg lay. Briefly, gravid adults were plated on OP50 K-agar plates and allowed to lay eggs for 4 hours at 15°C. Adults were then washed off the plates with K-medium (Boyd et al., 2012) leaving unhatched eggs behind; they were incubated at 15°C for 23 hours allowing eggs to hatch. At 23 hours post egg lay, L1 larvae were washed off plates, counted in liquid droplets, and plated at approximately 100 worms per plate on control or drug-treated 6 cm OP50-seeded K-agar plates. For growth assays the control plates (0 μM) were treated with the chemotherapeutic drug diluent 9 mg/ml sodium chloride, pH 3.6. Worms were incubated on the specified treatment plates for 3 days and subsequently imaged using a Leica M165 FC dissecting microscope. The Image J WormSizer plugin (Moore et al., 2013) was used to measure length and volume of each worm. Worm gonad images for length and -1 oocyte area measurements were taken of young adult animals aged 24 hours post mid-L4 under a 20X objective on a Leica DMI8 and measured manually in FIJI.

### DAPI Staining

DAPI staining was done by modifying standard protocols (Francis and Nayack, 2000), with the cold methanol fixation done for a shorter time (2.5 min) and the concentration of DAPI higher at 1 μg/ml in 0.01% Tween in PBS in the dark for 5 min, washed once with 0.1% Tween in PBS. Samples were briefly stored at 4°C in 75% glycerol and imaged directly in glycerol solution on a Leica DMI8 with an xLIGHT V3 confocal spinning disk head (89 North) with a ×63 Plan-Apochromat (1.4 NA) objective and an ORCAFusion GenIII sCMOS camera (Hamamatsu Photonics) controlled by microManager (Edelstein et al., 2010). DAPI was excited with a 405 nm laser. Worms were mounted on agar pads in 75% glycerol.

### Proliferative Zone and DAPI Body Measurements

Images of DAPI-stained young adult worm gonads (above) were analyzed for proliferative zone length and number of mitotic figures. Measurements were made in FIJI from the distal end of the gonad to the transition zone, which is the distal-most row of germ cells with more than one crescent-shaped nucleus. Mitotic figures were counted manually as metaphase or anaphase DAPI bodies. Observations of 0 mitotic figures were counted in the analysis. In the proximal gonad, the -1 oocyte (when visible) was imaged and DAPI bodies in the nucleus were counted.

### Embryo in Eggshell Measurements

Gravid adult hermaphrodites were cut with hypodermic needles in depression slides in M9 to release embryos, and embryos were mounted on agar pads in M9, coverslipped, and imaged on the Leica confocal system described above. Cross-sectional area was measured in FIJI in a central plane of focus.

### Reproduction Measurements

Diploid and tetraploid *C. elegans* populations were synchronized via egg lay as described above. Egg plates were incubated at 15°C for approximately 3 days to allow larvae to develop to L4 stage. L4 stage worms were then washed off the plates, counted by liquid droplet, and plated at approximately 100 worms per plate to control or drug-treated 6 cm OP50-seeded K-agar plates (freshly-made as described above on the same day worms reached L4 stage). The worms were incubated on these plates at 15°C for 24 hours.

To measure brood size and any reproductive defects brought about by ploidy differences and/or drug exposure, worms were picked and individually housed on normal OP50 plates (5 worms per group with experiments performed in triplicate). Each worm was transferred to a new plate every 48 hours. The eggs on the plates were allowed to hatch for 48 hours and then counted for viability (hatched larvae vs. unhatched eggs). This process was continued for the duration of the original worm’s reproduction.

To measure relative apoptosis in the germline, we adapted a previously published protocol for acridine orange staining (Lant and Derry, 2013). Briefly, after 24 hours of drug-exposure as described above, approximately 50 worms were picked to a normal OP50 plate and allowed to recover for two days. At this point 0.5 mL of 75 μg/mL acridine orange (abbreviated AO, Invitrogen, Fisher Scientific, Waltham, MA) solution was added dropwise and spread to coat the plate. Worms were incubated in the stain for 1 hour then promptly picked to a fresh OP50 plate and allowed to recover (de-stain) for at least 3 hours. 15-25 stained worms were then mounted to a 2% agarose pad on a microscope slide, paralyzed using 2.5 mM levamisole and imaged using a Leica DMI-8 Thunder Imager inverted microscope (ex. 470 nm/em. 510 nm). The worms were examined and scored for apoptotic bodies in the germline.

### Lifespan Studies

Diploid and tetraploid *C. elegans* populations were synchronized via egg lay as described above. Egg plates were incubated at 15°C until larvae had developed to L4 stage (approximately 3 days). At L4 stage, 40 worms for each group were picked to new OP50 plates and monitored daily until natural death. Worms were considered dead upon no response to 3 harsh touches with a platinum wire. During reproduction, they were transferred every three days to avoid confusion with any adult offspring.

### Mitochondrial Respiration

Oxygen consumption rates (OCR) were collected as a measure of mitochondrial respiration using Resipher by Lucid Scientific, Inc. The Resipher system consists of a Resipher 32X device for continuous OCR measurement, sensing lids for use on standard 96-well plates, and a central hub computer for real time data streaming. Raw oxygen concentration gradient of the liquid culture media is collected in real time as well as temperature and humidity conditions. Oxygen flux calculations are made continuously after a 2-hour initial equilibration period during which the media oxygen concentration gradient is established. The Resipher instrument allows for real time OCR measurement in living C. elegans in liquid culture. For these experiments, wells were seeded with age-synchronized L4-stage C. elegans at a density of 20 worms per well with complete K+ media, OP50 E. coli as a food source, and either vehicle (DMSO) or the ATP synthase inhibitor DCCD at 5 or 20 μM. Media blanks including E. coli were also included in all experiments. Worms were dosed in quadruplicate and oxygen consumption was collected by the Resipher continuously for 24 hours. Lethality was assessed visually and manually (worms were prodded with a platinum wire) following the experiments. Experiments were repeated a total of five times and data pooled.

### Transcriptional Analysis with RNA-seq

To determine the transcriptional changes underlying the phenotypic differences between tetraploid and diploid worms, whole-genome transcriptomic analysis was done by RNA-seq. In each group, 3,000 worms were pooled in each replicate for three biological replicates. L1 stage worm exposures were carried out to vehicle, 5 μM cisplatin, and 5 μM doxorubicin for 48-h as described earlier, except on large 10 cm plates to accommodate larger numbers of worms. Following exposure, worms were washed off plates into 15 ml centrifuge tubes and washed three times to remove bacteria. Worms were transferred to 1.7 mL Axygen tubes, allowed to gravity settle and as much liquid as possible removed from the tube. The worms were then suspended in 1 ml RLT buffer (Qiagen RNeasy kit) containing β-mercaptoethanol and flash-frozen in liquid nitrogen. After all replicates were complete and frozen, the samples were thawed and homogenized using a Bullet Blender at power setting 8 with 0.5 mm Zirconia/Silica beads. Samples were homogenized in five consecutive cycles with 30 sec homogenization, 1 min cooldown on ice until complete homogenization was observed under a dissecting microscope. The homogenate was then transferred to a new microcentrifuge tube and incubated at room temperature for 5 minutes to promote dissociation of nucleoprotein complexes. Next, homogenates were centrifuged at 12,000x g for 15 min at 4°C and the supernatant (approx. 600 uL) was mixed with an equal volume of 70% ethanol and loaded onto an RNA purification column (RNeasy, Qiagen). The purification was then carried out according to manufacturer’s instructions, including the optional on-column digestion with DNaseI to remove genomic DNA contamination. RNA quality check (yield, purity, and integrity), cDNA library construction, and Illumina sequencing were performed by NovoGene (Beijing).

Raw data of fastq format were processed by Novogene to remove adapter reads and low-quality reads. The clean paired-end reads were then mapped to the reference genome (wbcel235) using Hisat2 v2.0.5. Mapped reads were assembled by StringTie (v1.3.3b) and featureCounts v1.5.0-p3 was used to count the reads numbers mapped to each gene. Finally, the FPKM of each gene was calculated based on gene length and reads counts. Differential expression was performed using the DESeq2 R package (1.20.0). Genes were considered as differentially expressed if they had an absolute log_2_fold change of > 0.5 and q (adjusted p-value) < 0.05. Gene Set Enrichment Analysis was performed using the easy Visualization and Inference Toolbox for Transcriptome Analysis (eVITTA), which been validated in *C. elegans* datasets (Cheng et al., 2021) and uses up-to-date and species-specific databases for gene ontology. Pathways were considered to be enriched if the q (adjusted p-value) < 0.05. Raw RNA-seq data have been deposited in the National Center for Biotechnology Information’s GEO and are accessible through GEO series accession number GSE232747.

### Statistical Analyses

GraphPad Prism was used to perform statistical analysis. For all analyses, one- or two-way ANOVA tests were used, as appropriate, to determine the effect of ploidy and/or ploidy vs. treatment. Survival curves were generated using GraphPad Prism and statistical significance was determined using the Log-rank (Mantel-Cox) test. For RNA-seq data, differentially expressed genes were considered to be significant if they had a log_2_ fold change greater than 0.5 and an FDR-adjusted p-value less than 0.05. In eVITTA GSEA pathway analysis, enrichment was considered significant if the p and q values were less than 0.05.

## RESULTS

### Tetraploid *C. elegans* are larger and shorter-lived compared to diploid worms

After establishing stable tetraploid lines (Fig. 1a-b), we first evaluated the size of the animals relative to the genotype-matched diploid strain. The tetraploid worms were approximately 30% longer at young adulthood: the average length of Day 1 adult tetraploid worms was 1,433 μm while diploid was 1,119 μm. (Fig. 1c). This accords with previously published results (Clarke et al., 2018). We also observed that the lifespan of tetraploid worms was significantly reduced compared to diploid worms (Fig. 1d, individual traces shown in Supplemental Data Fig S1). The combined curves from all four replicates show a significant (p<0.001) reduction in lifespan from 20 to 17 days (Fig. 1d).

**Figure 1.**
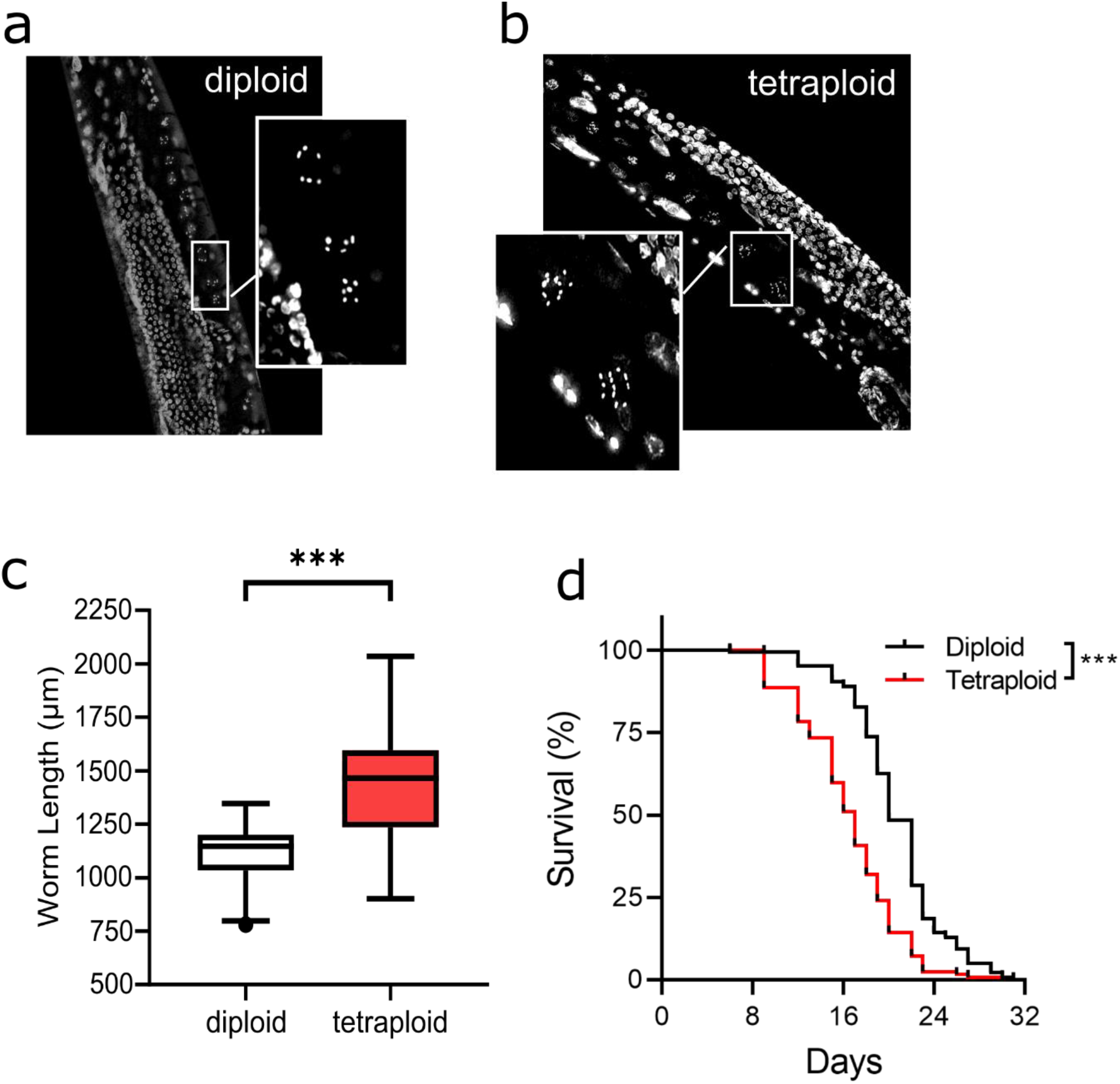
Tetraploid worms are longer and shorter-lived than diploid worms. <u>Panels a and b</u>, fluorescence image of DAPI-stained diploid and tetraploid worm mid-bodies, respectively. Insets highlight the two most mature unfertilized oocytes, during which stage the chromosomes condense into DAPI bodies that can be counted. Hermaphrodite diploid worms have 5 autosomes and 1 X-chromosome for a total of 6 DAPI bodies, while tetraploid worms have double (12). <u>Panel c</u>, length of worms was determined using brightfield microscopy images and the WormSizer plugin on ImageJ. Data are compiled from >8 biological replicates and >150 worms per ploidy group. Asterisks represent p<0.001, Student’s unpaired t-test. <u>Panel d</u>, lifespan curves for diploid and tetraploid worms are combined from four biological replicates (total N=142 (diploid) and N=128 (tetraploid); for individual traces, see Supplemental Fig S1). Asterisks represent significantly different survival curves p<0.001, Mantel-Cox Log-rank test.

### Despite a larger body size, tetraploid worms do not respire more and are not more sensitive to inhibition of ATP synthase

We anticipated that due to the larger size of tetraploid worms, they may require a higher rate of mitochondrial respiration. To test this, we measured the oxygen consumption rate beginning at the L4 stage for 24 hours (Fig. 2a). During this stage of development into adulthood, the oxygen consumption increased throughout the 24-h period. However, we discovered there is not a significant difference in tetraploid *C. elegans* respiration compared to diploid animals (Fig. 2b). Respiration was similar at all timepoints spanning L4 to young adult stages, and respiration of both strains was similarly inhibited by the ATP synthase inhibitor DCCD. Inhibition of ATP synthase by DCCD allows for the indirect measurement of oxygen consumption coupled to ATP production (Luz et al., 2015). We also evaluated survival following the respiration measurements and observed that both strains were sensitive to lethality from ATP synthase inhibition by DCCD to a similar degree (Fig. 2c). We observed high variability in lethality at the 5 μM dose, but no consistent differences between the ploidy groups. Taken together these results indicate that while tetraploid worms are approximately 30% larger, they are not increasing mitochondrial respiration to compensate for their larger size.

**Figure 2.**
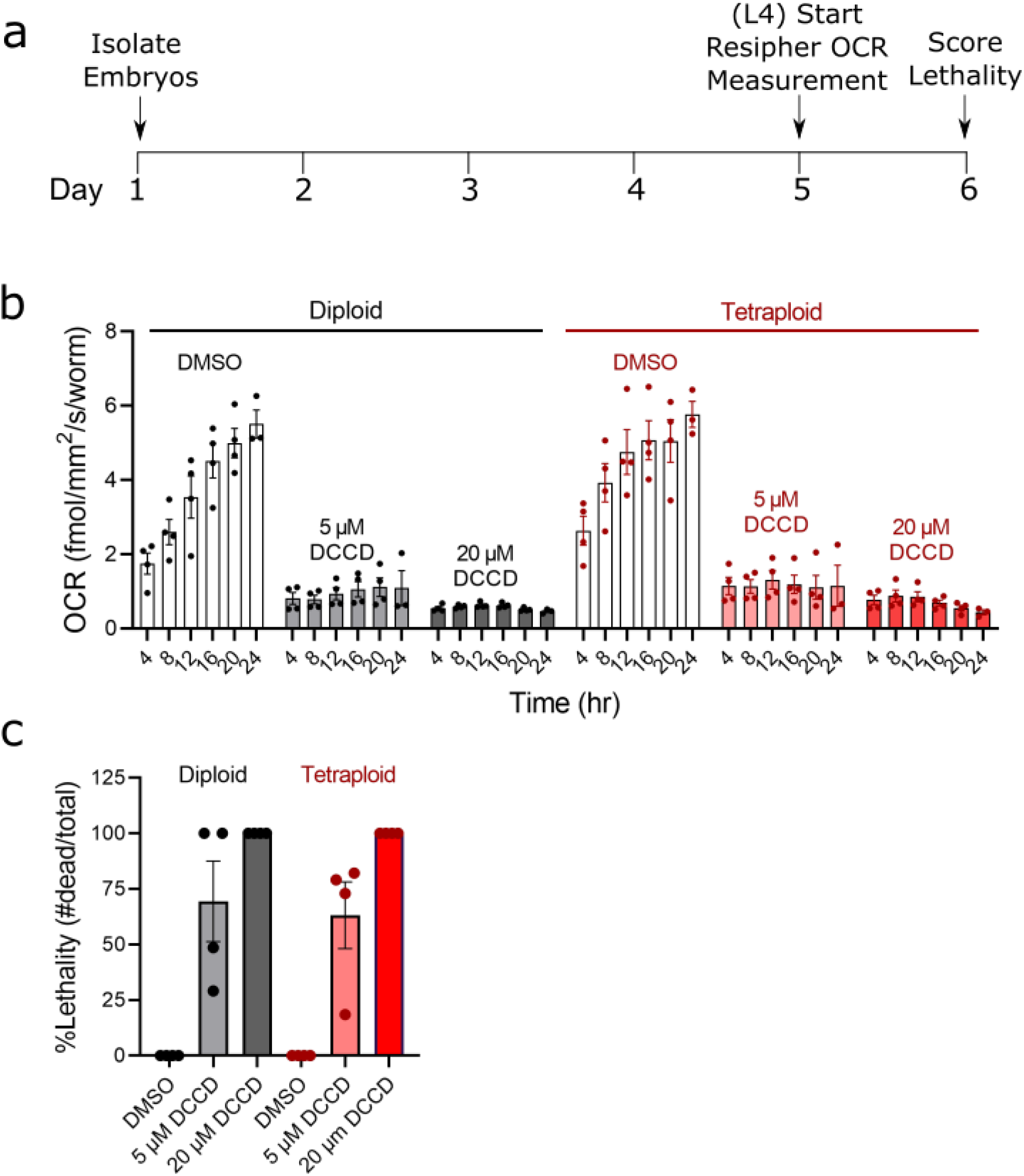
Despite larger size, polyploid worms do not have higher respiration than diploid worms. <u>Panel a</u>, timeline of dosing and measurements. For experiments, age-matched embryos were obtained via egg lay with gravid adults, then raised to the L4 stage at 15°C. Worms were then picked into 96-well plates for Resipher (Lucid Scientific) analysis at a density of 20 worms/well in a volume of 100 μL containing OP50 *E. coli* bacteria for food and vehicle (0.5% DMSO), 5 μM, or 20 μM DCCD (ATP synthase inhibitor). OCR was measured continuously for 24h, following which lethality was visually assessed under the microscope after a harsh touch with a platinum wire. <u>Panel b</u> shows OCR at 4h time intervals throughout the measurement, normalized per worm. Each data point shown is the average of 4 technical replicates. Experiments were carried out in 4 biological replicates. Bars and error bars show the mean and SEM, respectively. No significant differences were observed between ploidy groups in a two-way ANOVA with Bonferroni-adjusted post testing for individual comparisons. <u>Panel c</u> shows the lethality following DCCD exposure in the Resipher plate. No significant differences were observed between ploidy groups.

### Diploid and polyploid worms do not differentially express genes related to mitochondrial function or energy metabolism

We performed mRNA sequencing to examine transcriptional differences between the ploidies. Pathway analysis revealed the most up- and down-regulated pathways (Table 1). Interestingly, many pathways involved in DNA and RNA synthesis and handling and nuclear division were downregulated in tetraploid worms, while upregulated pathways included signaling pathways, unfolded protein response, major sperm protein, and structural pathways involving collagen.

**Table 1.**
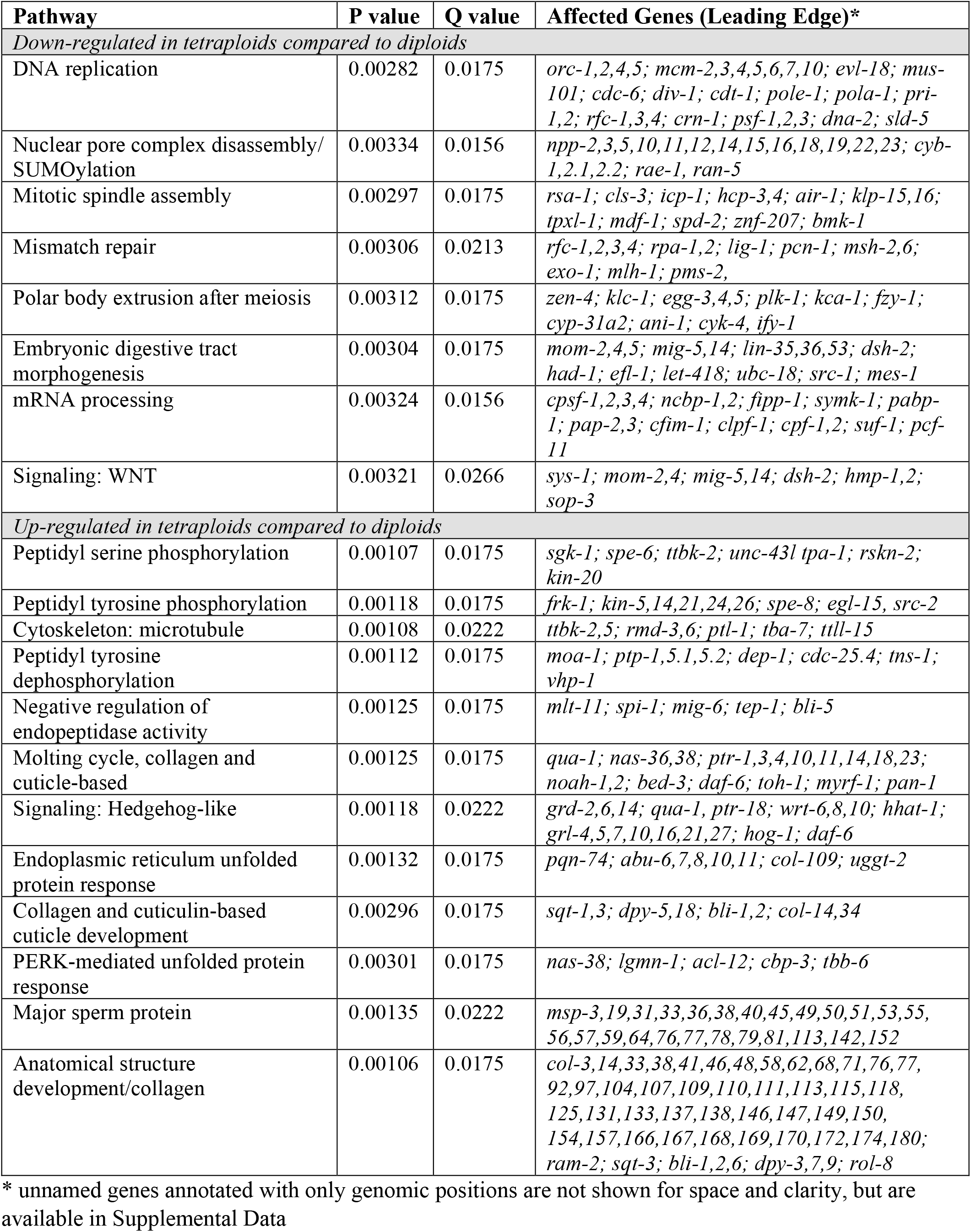
Significantly enriched pathways in tetraploid vs. diploid worms.

In agreement with our findings that tetraploid worms do not respire at higher rates than diploid worms, we did not see significant enrichment in genes related to mitochondrial function between diploid and tetraploid worms, or enrichment of other energy metabolism pathways such as glycolysis or fatty acid oxidation. We did see a trending significance of sphingolipid pathways, which have been previously implicated in polyploid cancer cells (Lu et al., 2021). In contrast to studies in cancer cells, we observed no change or downregulation of the ceramides in our study (Supplemental Fig. S2). Together, these results suggest that there is not a dramatic difference in energy requirements between diploid and tetraploid worms.

### Polyploidy reduces reproductive output, proliferative germline size, mitotic index, and robustness of chromosomal segregation

We evaluated the reproductive fitness of tetraploid worms compared to diploids by measuring total brood size and reproductive defects (dead unhatched embryos and sterility, Fig. 3a). We observed a significant reduction: approximately 70%, in viable offspring, ∼100 offspring per tetraploid worm compared to ∼300 for diploid worms (Fig. 3b), which agrees with previous findings (Madl and Herman, 1979). Additionally, the percentage of unhatched embryos was significantly increased in tetraploid worms, >13%, compared to just ∼3% in the diploids (Fig. 3c). While tetraploid worms lay fewer eggs in total, this alone does not account for the decrease in viable offspring; tetraploid worms also experience increased frequency of nonviable fertilized eggs.

**Figure 3.**
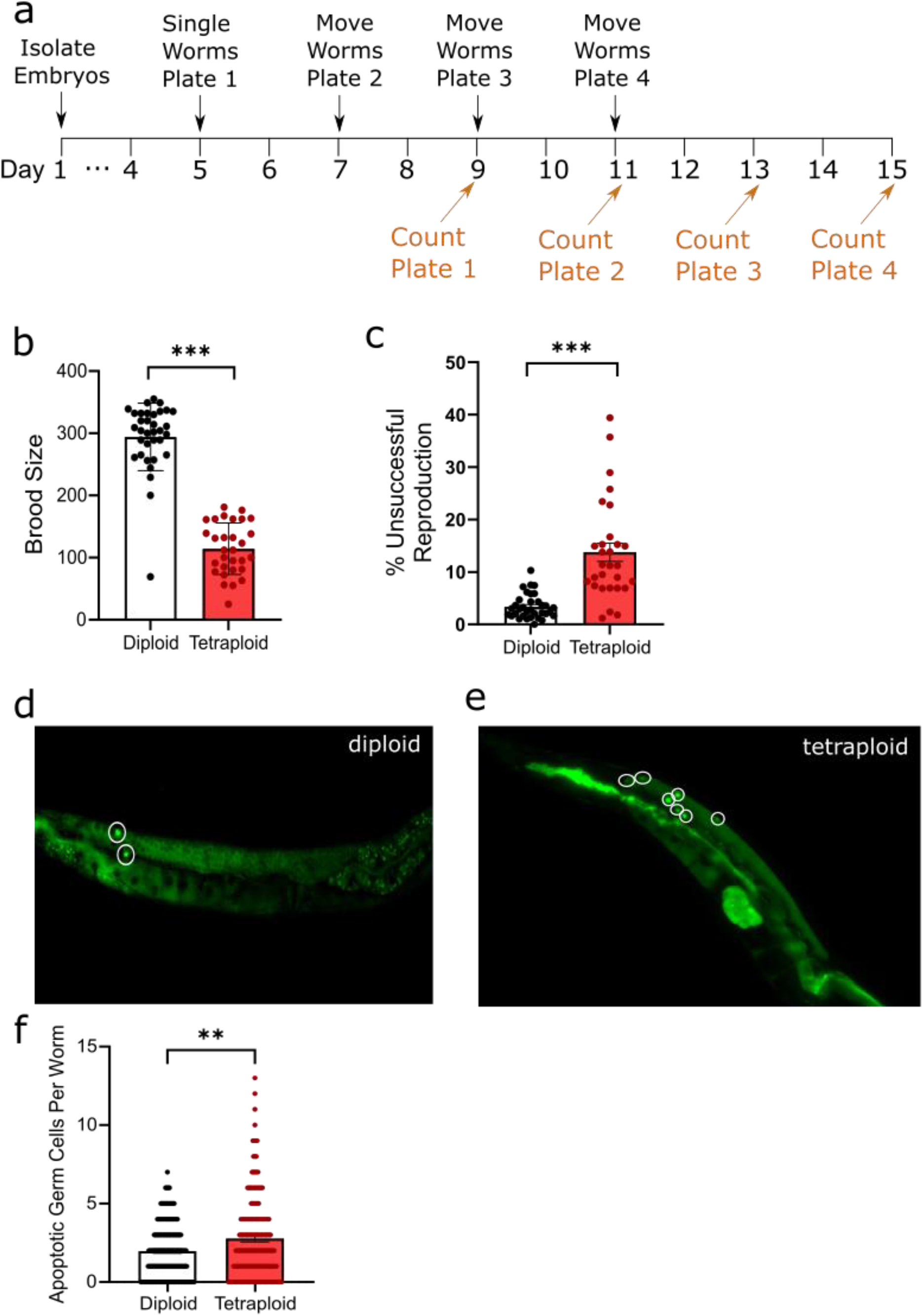
Polyploidy reduces reproductive fitness. <u>Panel a</u>, timeline of reproduction measurements. For experiments, age-matched embryos were obtained by an egg lay with gravid adults on day 1. On day five worms were singled (five per group) to individual plates and allowed to lay eggs. They were moved to new plates every 48h a total of three times, and the eggs were allowed to hatch for an additional 48h before the plates were counted. Live embryos were counted and summed for brood size; eggs unhatched after 48h were considered dead embryos. Worms that did not produce any eggs at all were counted as sterile. <u>Panel b</u>, brood size for diploid and tetraploid worms compiled from 5-6 biological replicates. Each data point represents the summed total living offspring from a single worm. <u>Panel c,</u> percent unsuccessful reproduction events calculated from the same 5-6 biological replicates. Unhatched eggs (dead embryos) were counted and divided by the total eggs laid (viable larvae + unhatched eggs). Each data point represents the percent of unhatched eggs from a single worm. <u>Panels d-e</u>, representative images of acridine orange staining. On day 5 worms not picked for reproduction experiments were exposed to 75 μg/mL AO for one hour, then removed and allowed to recover for 3 hours prior to imaging. <u>Panel f</u>, quantification of AO positive cells per worm in diploid and tetraploid worms. *** represent p<0.001, ** represent p=0.002, Student’s unpaired t-test

Dead embryos are suspected to be aneuploid (having inherited the wrong number of chromosomes.) We therefore examined the embryos and went upstream to examine the germline for defects. Apparently healthy tetraploid embryos are ∼1.50x larger in cross-sectional area than diploid embryos (Fig. 4d), implying the size difference that persists throughout life is evident from embryogenesis. The -1 oocyte (the oocyte next to the spermatheca, which is the most mature oocyte in the gonad arm and is about to be ovulated) of tetraploids is ∼1.33x larger (Fig. 4c).

**Figure 4.**
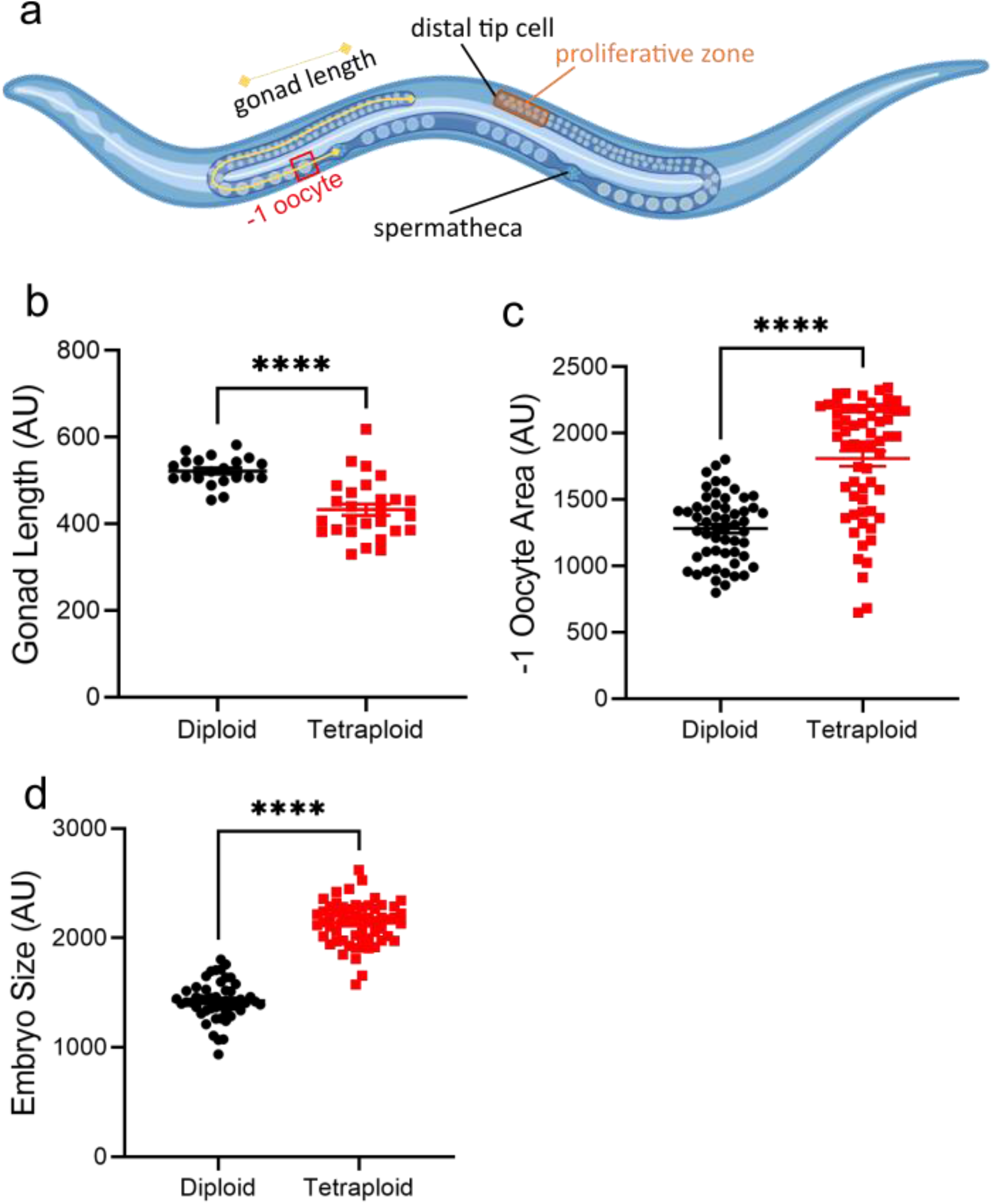
Tetraploid worms produce larger offspring with a shorter total germline length. <u>Panel a</u>, schematic representation of the two symmetrical U-shaped arms of the *C. elegans* hermaphrodite germline that join at a common uterus. Key features are annotated including the distal tip cell, spermatheca, -1 oocyte, and proliferative zone. Figure created using Biorender.com. <u>Panel b</u>, length of gonad measured from distal tip cell to the spermatheca using image analysis of DAPI-stained fixed adult worms. <u>Panel c</u>, area of -1 oocyte. <u>Panel d</u>, area of embryos. ****, p<0.0001, Student’s unpaired t-test.

Since physiological apoptosis in the germ line serves a nurse-cell-like function in oogenesis, we hypothesized that larger oocytes might be correlated with more apoptosis. We used acridine orange staining to evaluate any differences in the number of apoptotic germ cells between the groups (Fig. 3d-f). Our experiments showed a significant increase in germline apoptosis in tetraploid *C. elegans* compared to the diploids. There is evidence that binucleate germ cells are removed by physiological apoptosis in *C. elegans* diploids (Raiders et al., 2018), so it is possible that aneuploidy in the meiotic germline is responsible for the elevated rates of apoptosis we observe. These hypotheses—that aneuploidy causes elevated apoptosis and elevated apoptosis increases oocyte size—are not mutually exclusive.

While we interpret decreased brood size and increased embryonic death as evidence for aneuploidy, these fertility defects may be attributable to other causes. We next assayed aneuploidy directly in the -1 oocyte by DAPI staining, in which DAPI bodies can be individually counted. Normalizing to the expected number of bivalents (six for diploids and 12 for tetraploids), the tetraploids (N=30) display significantly more aneuploidy than diploids (N=39, mean difference from expected is 15.8% vs. 6.5% for diploids, Welch’s t-test, t=3.4206, df=49.924, p-value=0.001254). As expected, there is a higher frequency of aneuploidy in tetraploid gametogenesis.

Aneuploidy is also reflected in the frequency of male offspring because the male sex is specified in *C. elegans* by the presence of only one X-chromosome (hermaphrodites have two X-chromosomes). We observed a 10-fold increased rate of males in the tetraploid lines (mean frequency 1.43%, ± 0.33% standard error of the mean, compiled counts from 14 independent plates with n>100 adults/plate) compared to diploid wild-type animals, which have a frequency of males between 0.1-0.3% (Quevarec et al., 2022, Hodgkin, 1983, Chasnov and Chow, 2002). Our observed higher incidence of males provides additional support for the higher rate of aneuploidy in the tetraploid lines.

Going further upstream in the gonad, we found that tetraploid gonads were nearly 20% shorter than diploid gonads (single gonad arm measured from distal tip to spermatheca, Fig. 4a-b). The proliferative zones (between the distal tip and the crescent-shaped pachytene nuclei in the meiotic region) were not significantly different in length, and the incidence of germ cell division also did not differ (data not shown). Taken together, tetraploids have roughly the same amount of germ cell proliferation in the same size gonad region, followed by significantly more apoptosis in a germline making larger eggs and embryos. This leads to the gonad length being diminished.

### Polyploidy modestly protects against drug-induced growth delay

After evaluating the physiological differences between tetraploid and diploid worms, we wanted to test their resilience in the face of stressors. At the simplest level, polyploidy may protect cells during stress by buffering DNA damage with more functional copies of each gene. Therefore, we tested the impact of tetraploidy on sensitivity to DNA damage-inducing chemotherapeutic drugs cisplatin and doxorubicin.

Cisplatin and doxorubicin both intercalate into DNA and cause DNA lesions through alkylating nucleotides, and in the case of cisplatin, causing DNA crosslinks. We exposed diploid and tetraploid worms to these agents by adding the drugs to agar plates and placing the worms on those plates for 72 hours beginning at the L1 stage. After 72h, we imaged the worms and measured their length (Fig. 5a).

**Figure 5.**
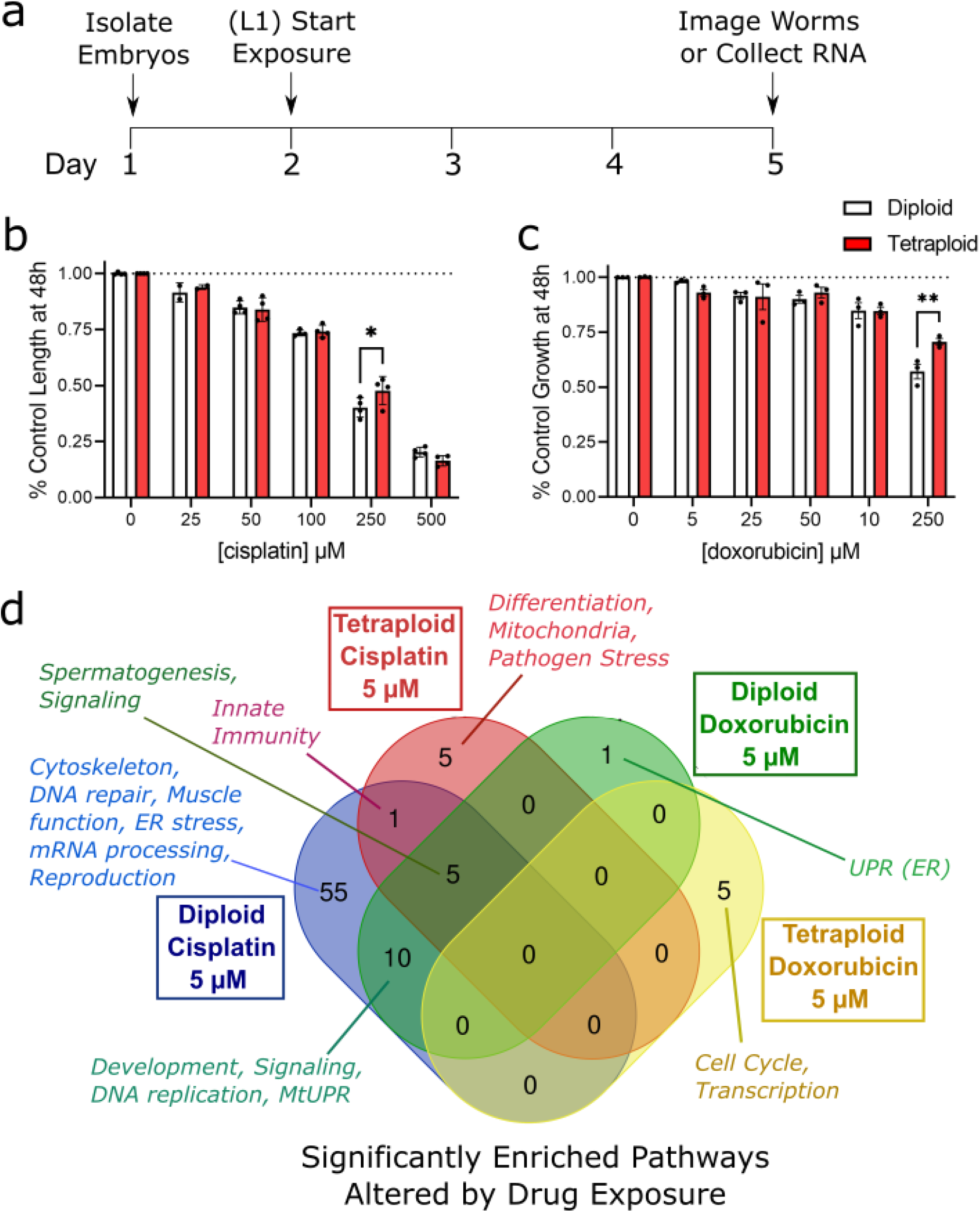
Tetraploid worms are modestly protected from growth delay at high doses of chemotherapeutics. <u>Panel a</u>, timeline of dosing and measurements. For experiments, age-matched embryos were obtained by an egg lay with gravid adults on day 1. Eggs were allowed to hatch for 24 h and larvae were counted and moved to plates containing cisplatin or doxorubicin. The worms developed at 15°C on drug plates for 72 h (until the L4 stage), at which point they were either imaged for size determination or frozen down for RNA isolation. <u>Panels b-c</u> show dose-dependent effects on growth for cisplatin and doxorubicin, respectively. Growth data is shown as relative size compared to the control (due to size differences between diploid and tetraploid control worms). Asterisks represent significance in a two-way ANOVA (ploidy vs. dose) with Bonferroni-corrected p<0.05 in multiple comparison post testing. <u>Panel d</u>, Gene Set Enrichment Analysis using eVITTA revealed top significant pathways altered in drug-exposed tetraploid worms vs. diploid worms with FDR-adjusted p<0.05.

Both drugs caused a dose-dependent decrease in growth in both ploidy groups with cisplatin being more growth-inhibiting than doxorubicin (Figs 5b-c); at the highest dose, cisplatin caused L1 larval arrest for both diploids and tetraploids, so that dose was discarded in further analysis. For the highest dose of both drugs that caused growth inhibition without developmental arrest, tetraploidy was modestly protective.

For cisplatin, a two-way ANOVA revealed a significant effect of dose (p<0.0001) and a significant interaction between ploidy and dose (p = 0.0465). Multiple comparison analysis revealed significant protection in tetraploid worms (p = 0.0193) at the 250 μM dose (Fig. 5b). In the case of doxorubicin, there was a significant effect of dose (p<0.0001) and a significant interaction term (p=0.0278) in the two-way ANOVA. Multiple comparison analysis revealed protection of tetraploid worms at the highest dose of 500 μM (p=0.0061, Fig. 5c). Together, these results show that tetraploid worms are somewhat protected from growth inhibition at doses that cause roughly 50% growth delay in diploid strains.

### Tetraploid worms respond less robustly at the transcriptional level than diploid worms to low doses of chemotherapeutics

Following up on the growth experiments, we wanted to characterize responses to chemotherapeutics using mRNA sequencing. Based on our growth delay experiments, we selected doses of each drug that would not cause dramatic (>10%) growth delay (5 μM) to avoid any confounding effects of developmental differences on gene expression. Overall, we found that the tetraploid worms had fewer differentially expressed genes related to the chemotherapeutic exposure than the diploid worms, which is apparent in the Principal Component Analysis plot (Supplemental Fig. S3). In diploid animals, both DNA-damaging chemotherapeutics triggered differential expression of pathways related to development, signaling, DNA replication, and the mitochondrial UPR, while cisplatin alone triggered many transcriptional changes in diploid animals including DNA repair, cytoskeleton, muscle function, ER stress and proteostasis, and mRNA processing (Fig. 5d). Full pathway analysis is given in Supplemental Materials.

### Polyploidy does not protect worms from reproductive toxicity from cisplatin and doxorubicin

In *C. elegans*, the germline is the most mitotically active post-embryonic tissue. While the number of somatic cells will not quite double between hatching and adulthood, the number of germ cells increases by a factor of over 1000. Since both cisplatin and doxorubicin inhibit DNA replication, we hypothesized that the effects of these drugs on the germline and the soma of *C. elegans* might be different.

While the growth delay we observed may reflect drug toxicity inhibiting mitotic cell divisions during development, we also wanted to test how these drugs impact *C. elegans* reproduction. To test this, we exposed worms for 24 h bridging from the L4 larval stage to young adulthood to target a vulnerable period of germline expansion and measured total brood size and reproductive defects in drug-exposed worms (Fig. 6a).

**Figure 6.**
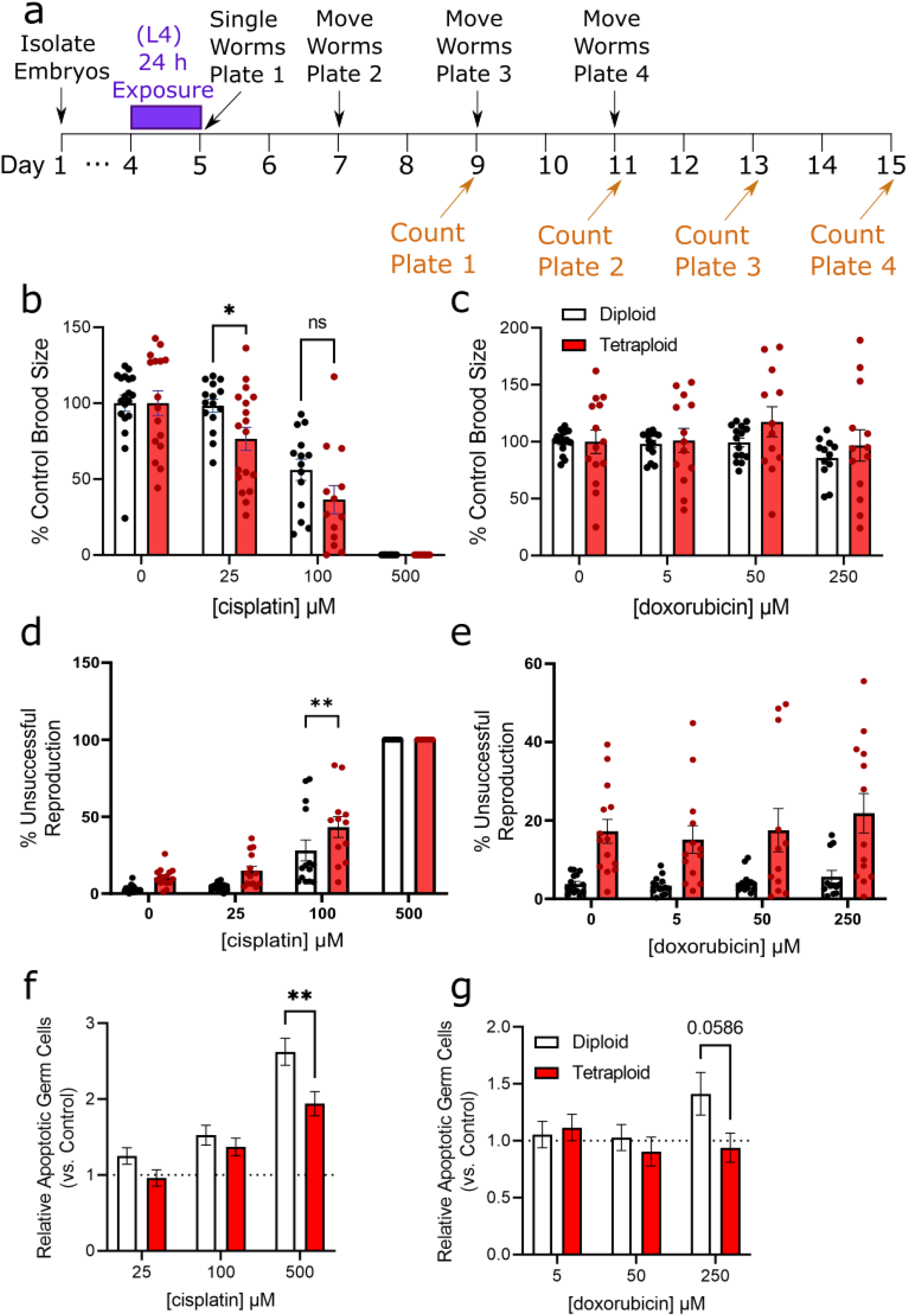
Tetraploid worms are more sensitive to cisplatin and doxorubicin-induced reproductive toxicity. <u>Panel a</u>, timeline of dosing and reproduction measurements. For experiments, age-matched embryos were obtained by an egg lay with gravid adults on day 1, and transferred to agar plates containing cisplatin or doxorubicin for 24h at the L4 stage at 15°C. Following exposure, worms were singled (5 per group) to individual plates and allowed to lay eggs. They were moved to new plates every 48h a total of three times, and the eggs were allowed to hatch for an additional 48h before the plates were counted. Live embryos were counted and summed for brood size; eggs unhatched after 48h were considered dead embryos. Worms that did not produce any eggs at all were counted as sterile. Worms that crawled off the plate were censored. <u>Panels b-c</u>, brood size for diploid and tetraploid worms at varying doses of cisplatin and doxorubicin. Each data point represents the summed total living offspring from a single worm, compiled from 3 biological replicates (total N=13-18 hermaphrodites/group). <u>Panels d-e</u>, frequency of (aneuploid) dead embryos and sterility in cisplatin and doxorubicin-exposed worms, respectively. <u>Panels f-g</u>, relative number of apoptotic germ cells in cisplatin and doxorubicin-exposed worms relative to control. Asterisks represent significance in a two-way ANOVA (ploidy vs. dose) with Bonferroni-corrected p<0.05 in multiple comparison post testing.

Exposure to cisplatin, but not doxorubicin, caused a dose-dependent decrease in brood size for both diploid and tetraploid worms (Figs. 6b-c). A two-way ANOVA for cisplatin revealed significant effects of ploidy, dose, and the interaction (p<0.0001 for all terms). Bonferroni-corrected multiple comparison analysis revealed significant reduction in brood size at 100 and 500 μM for both ploidy groups (p<0.001 in all cases), and only for tetraploid worms at 25 μM cisplatin (p=0.0183). For doxorubicin, two-way ANOVA analysis revealed a significant effect of ploidy (p<0.0001), and a significant effect of dose (p=0.0488), but no significant interaction. Furthermore, there was a significant reduction in brood size at the highest dose 250 μM in diploids only (p=0.0179).

In addition to counting live offspring, we also counted unhatched eggs on the plates and noted two diploid and three tetraploid sterile worms in the cisplatin exposed groups (Figs. 6d-e). Two-way ANOVA analysis for cisplatin revealed a significant effect of ploidy and dose (p<0.001) but no significant interaction (p=0.13). We observed that in the cisplatin 100 μM dose, there was a higher rate of unsuccessful reproduction (dead embryos) in the exposed tetraploid worms compared to the diploid worms. There was a dose-dependent increase in aneuploidy (dead embryos) for both ploidies with cisplatin, causing the most dramatic effect at the 500 μM dose (Fig. 6d). There were no successful reproduction events among either diploids or tetraploids at this dose (i.e. none of the eggs laid hatched to larvae). For doxorubicin, two-way ANOVA was not significant for ploidy, dose, or interaction terms. Accordingly, we observed dramatically higher rates of unsuccessful reproduction in the tetraploid worms compared to the diploid worms but did not observe any significant dose-dependent increases in that rate (Fig. 6e).

Induction of DNA damage is known to increase germline apoptosis in diploid *C. elegans* (Schumacher et al., 2001), so we tested whether we would see differential induction of apoptosis between the ploidy groups. For cisplatin, two-way ANOVA revealed significant effects of ploidy and dose (p<0.001) but no significant interaction (p=0.14). For doxorubicin, there was no significant effect of ploidy or dose or interaction in the two-way ANOVA. Both drug treatments increased the relative number of apoptotic germ cells in diploid worms to a greater degree than in tetraploid worms, though the effect was only significant for cisplatin (Figure 6f and 6g).

Together, these results suggest that tetraploidy is not protective against DNA-damaging agents administered during reproductive adulthood, and that tetraploidy may predispose worms to aneuploidy under stressful conditions. Thus, DNA-intercalating agents may impose a higher cost on mitotically active polyploid cells.

## DISCUSSION

In this study, we used a recently published method to generate stable lines of tetraploid worms as an in vivo model of polyploidy. We then used this model to study the gene expression and physiological consequences of polyploidy, with an application of how stress from chemotherapeutic drugs affect polyploid vs. diploid organisms. Tetraploid animals have a shortened lifespan, longer bodies, shorter germ lines, and reduced brood size with modest protection from growth effects but not reproductive defects induced by DNA-damaging drugs cisplatin and doxorubicin. We report many transcriptional differences between diploid and tetraploid animals at baseline and after exposure to both drugs. After low doses of both drugs were administered, we saw a strong transcriptional response only in the diploid animals with very little differential gene expression in the exposed vs. unexposed tetraploid animals. Together, these findings suggest that short-term transition to tetraploidy in worms during stress may buffer them against environmental insults.

In the RNA-seq data comparing diploid to tetraploid worms without drug exposure, we observed at least 2-fold downregulation of all four of the *C. elegans* cyclin B family members that are orthologs of human cyclin B3, the *C. elegans* cyclin D1/D2 homolog *cyd-1*, and similar downregulation of all of the *C. elegans* cyclin-dependent kinases relative to diploids. Interestingly, this did not produce the severe developmental defects that have been observed in RNAi against these genes, perhaps due to differing levels of expression loss. We hypothesize that these genes are expressed at a lower level to permit synthesis of the extra set of chromosomes during developmental cell divisions, but precisely how the system is tuned to allow tetraploidy but prevent deleterious loss of these functions has not yet been investigated.

We observed a ∼30% increase in worm length in the tetraploid worms compared to diploid. This particular size increase seems to be common, as tetraploid yeast are ∼30-50% larger than diploid yeast (Otto, 2007), and mammalian cells have also been reported to increase in size by approximately ∼40% when they become tetraploid (Hau et al., 2006). This conserved increase in size is not universal, however. Fankhauser observed that higher ploidy levels in salamander tissues resulted in bigger but fewer cells, in order for organ size to remain the same. Similarly, while tetraploid mouse embryonic cells were increased ∼2-fold compared to diploid, the embryos themselves were not increased in size. There were less than half the number of cells in the tetraploid tissues as compared to diploids (Henery et al., 1992), suggesting a regulation of tissue size independent of cell size regulation. In liver with partial hepatectomy, hypertrophy of cells and increased ploidy accounts for a significant portion of the regenerated tissue mass (Miyaoka et al., 2012). The mechanisms relating ploidy to cell, organ, and organism size are still not well defined.

It is interesting that the mitochondrial function of the two strains was similar. One of the prevailing theories behind polyploid induction in cells is that it may increase the cell’s overall capacity for molecular functions of life. It is hypothesized that multiple copies of DNA allow for more mRNA and protein synthesis, making cells themselves larger in size and allowing for increased functionality. We see the increase in overall organism size, but no concomitant increase in mitochondrial respiration. It is possible that increased respiration may only be utilized on an “as-needed” basis. For example, in this study we did not examine the worms’ maximal respiratory and spare respiratory capacity, but this could be tested in future studies. However, previous studies in polyploid giant cancer cells showed very modest differences in respiration including maximal respiratory and spare capacity (Lu et al., 2021), suggesting that the increased ploidy does not necessarily increase mitochondrial capacity. In response to both chemotherapeutic drugs, the diploid animals activated the mitochondrial unfolded protein response (MtUPR) while the tetraploid animals did not. Activation of the mitochondrial UPR in breast cancer cells has previously been demonstrated to promote resistance to cisplatin (Chen et al., 2021). This may add a layer of complexity to the comparative sensitivity of diploid vs. tetraploid animals to cisplatin toxicity – while tetraploid animals may have other defenses, they fail to activate this protective mechanism in response to the drug treatments.

We observed rather modest protection from growth delay at medium- to high-dose exposure to DNA damaging agents. The protection was observed at doses that caused significant growth delay, but not arrest, in diploid animals. We observed complete L1 arrest at the highest dose of cisplatin in both ploidy groups, indicating this level of damage cannot be overcome even with more copies of the genome. A previous study showed that in liver and fibroblast cell lines that had been converted to stable tetraploidy, ionizing radiation caused a dramatic increase in the number of γ-H2AX foci in tetraploid cells compared to diploid cells, indicating increased DNA damage. This increase was far greater than the 2-fold increase in DNA content (Hau et al., 2006). We made a consistent finding, in that tetraploid germlines – the most mitotically active tissue that we have studied – are not protected from DNA damaging agents, and in fact are somewhat more fragile than diploid germlines. Our transcriptional data are suggestive of repressed DNA damage machinery in polyploids, a finding which is also supported by microarray experiments in tetraploid vs. diploid primary hepatocytes (Lu et al., 2007).

In contrast to the early developmental exposure, the late larval L4 exposure during expansion of the germline did not reveal protection of the tetraploid animals. The tetraploid animals had similar or greater increases in aneuploidy compared to the diploid worms when exposed to cisplatin. Previous studies in tetraploid hepatocytes (Duncan et al., 2012, Wilkinson et al., 2019), Drosophila (Fox et al., 2010, Schoenfelder et al., 2014), and polyploid cancer cells (Fujiwara et al., 2005, Was et al., 2022) have demonstrated that polyploid cells are more likely to give rise to aneuploid cells. Our findings support a similar or greater propensity to produce aneuploid embryos in tetraploid nematodes compared to diploid.

Altogether, the findings of this study give an unprecedented view into the physiological consequences of whole-animal tetraploidy in *C. elegans*, including the impacts on gene expression, metabolism, lifespan, reproduction, and response to chemotherapeutic exposures. Surprisingly, although tetraploidy is a natural stress-induced state in *C. elegans*, tetraploidy afforded only modest protection from DNA damaging agents during development. Further studies in this powerful model of polyploidy are needed to understand the biological impetus for polyploidization and ploidy reduction with natural stressors, and the genetic pathways that underlie those changes.

## Supporting information

Supplemental Spreadsheet - DEGs

Supplemental Spreadsheet - Pathways

## Acknowledgements

The authors would like to thank Dr. Mara Schvarzstein (Brooklyn College-CUNY, New York, NY) for her kind assistance in establishing tetraploid lines and Jason Pierce in the Lipidomics Core for his assistance with preparing samples for lipidomics. This work was supported in part by the American Cancer Society Institutional Research Grant #IRG-19-137-20, from the American Cancer Society, the Hollings Cancer Center at the Medical University of South Carolina, the National Institutes of Health R00-ES029552 (JHH), R35-GM147704 (KLG), T32-GM132055 (EAA and JBD), R25-GM113278 (HCG) and the Lipidomics Shared Resource, Hollings Cancer Center, Medical University of South Carolina (P30 CA138313 and P30 GM103339).

## Author Contributions

K.R.M., E.A.A., C.V.J., K.L.G., and J.H.H. conceived and designed the experiments. J.H.H. and C.V.J. conceived and designed the respiration, chemotherapeutic exposures, and transcriptomic study. J.H.H. and K.L.G. conceived and designed the reproduction experiments. K.R.M., E.A.A., H.C.G., K.A.S., X.L., C.M., B.S., J.B.D., K.L.G., and J.H.H. performed the experiments and analyzed the data. K.R.M. and E.A.A. wrote the first manuscript draft. All authors edited, read, and approved the final manuscript.

## Availability of materials and data

The datasets used and analyzed during the current study are available from the corresponding author upon reasonable request.

## Notes

### Competing Interest Statement

The authors have declared no competing interest.

