## Supplemental Figures for "The consequences of tetraploidy on *Caenorhabditis elegans* physiology and sensitivity to chemotherapeutics"

**TABLE OF CONTENTS**

**Supplementary Fig. S1. Individual lifespan traces ……………………………	3**

**Supplementary Fig. S2. LC/MS results of sphingolipid levels ……………….	4**

**Supplementary Fig. S3. Principal Component Analysis for RNA-seq data …	5**

**Supplementary References ……………………………………………………… 6**

**EasyGSEA pathway analysis …………………………………………………… excel file**

**Differentially expressed gene lists ……………………………………………… excel file**

**Figure S1. Individual traces for lifespan experiment.** Each trace represents a lifespan experiment from an independent biological replicate with n=30-40 animals per group. Lethality was determined by response to a harsh touch with a platinum wire. Animals that did not respond to three repeated touches were considered dead. Worms that crawled off the plate were censored in the analysis.

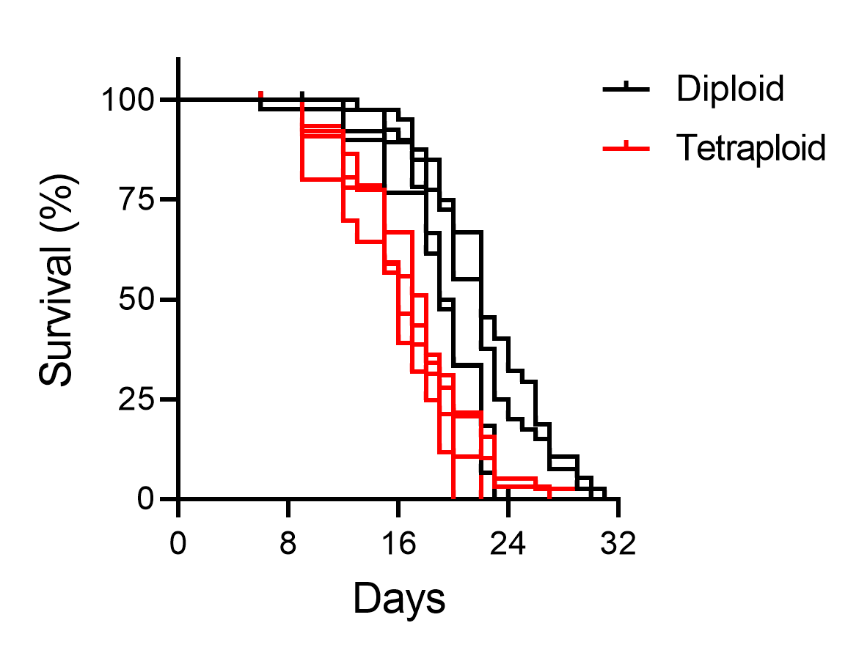

**Figure S2. LC/MS results of sphingolipid levels.** For experiments, L4-stage diploid and tetraploid nematodes were washed off plates, rinsed, and allowed to clear their guts, then centrifuged at 2200 x g for 2 min to pellet the worms. The worms were then resuspended in 1 mL Tissue Homogenization Buffer A (0.25M sucrose, 25 mM KCl, 50 mM Tris.HCl, 0.5 mM EDTA, pH 7.4) and lysed using a Q55 Sonicator (Qsonica LLC, Newtown, CT, USA). The worms were sonicated 10 times on ice, 40% power for 30 seconds followed by >2 min rest for each cycle. The protein content in the resulting lysate was determined using the Pierce BCA Assay (Thermo Scientific, Waltham, MA, USA). Protein content ranged from 2.6-4.3 mg/ml. Samples were then flash-frozen and stored at -80°C until they were submitted to the Medical University of South Carolina Lipidomics Core Facility, where the lipids were extracted with isopropanol:water:ethyl acetate (30:10:60 v:v:v). The individual lipid species were quantified by LC/MS as previously described*^1^*. Asterisks represent significance between diploid and tetraploid in two-way ANOVA with Bonferroni-corrected multiple comparisons: *, p<0.05; **, p<0.01; ****, p<0.0001.

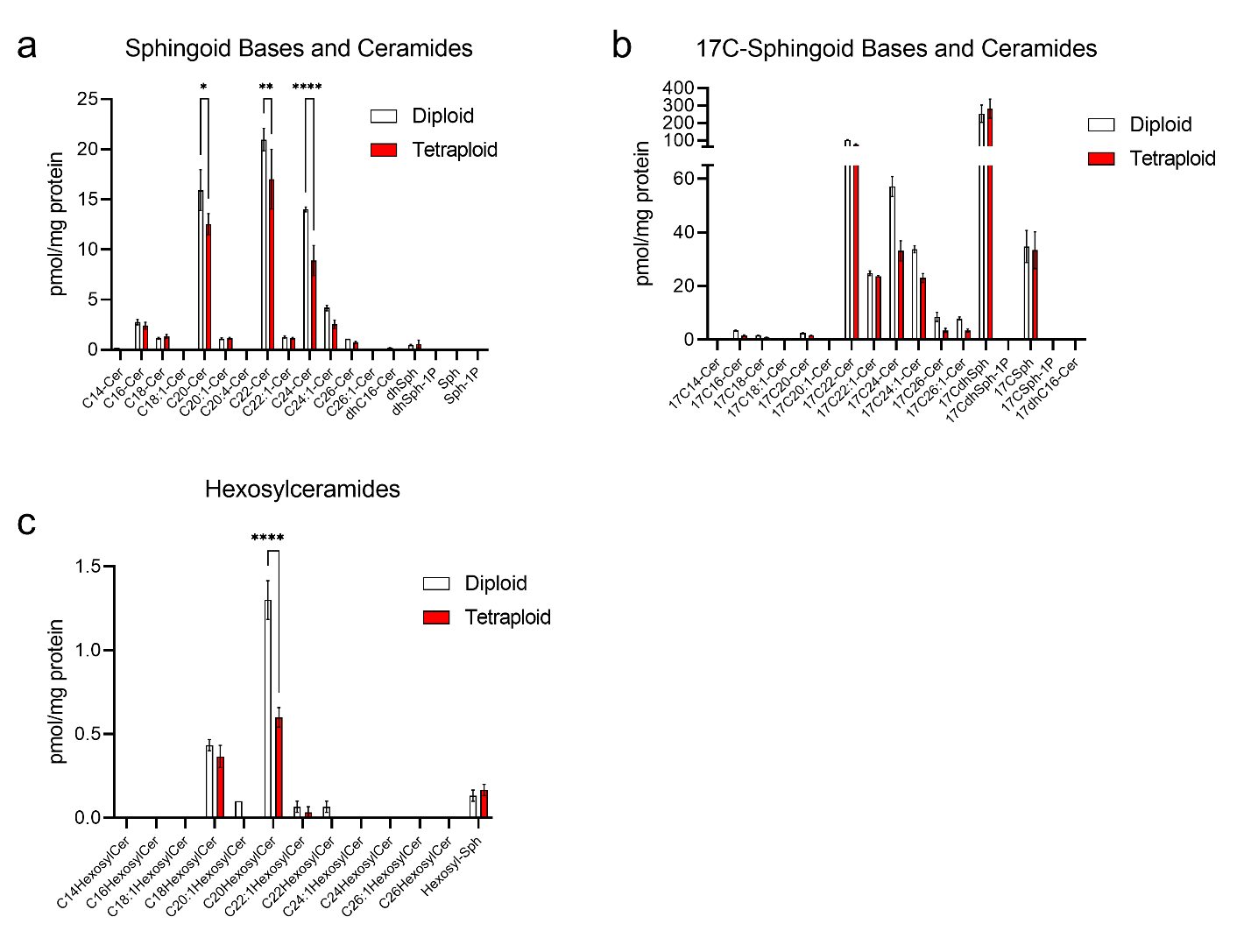

**Figure S3. Principal Component Analysis 2D plot.** For experiments, RNA was isolated from L4 stage worms exposed to vehicle (diploid and tetraploid, Dipl_CTRL and Tetra_CTRL), 5 µM cisplatin (Dipl_CIS and Tetra_CIS), or 5 µM doxorubicin (Dipl_DOX and Tetra_DOX). The numbers on the plot (1, 2, and 3) reflect the biological replicate for that data point. Principal component analysis was performed by Novogene (Sacramento, CA, USA) to check for inter-sample and inter-group variability.
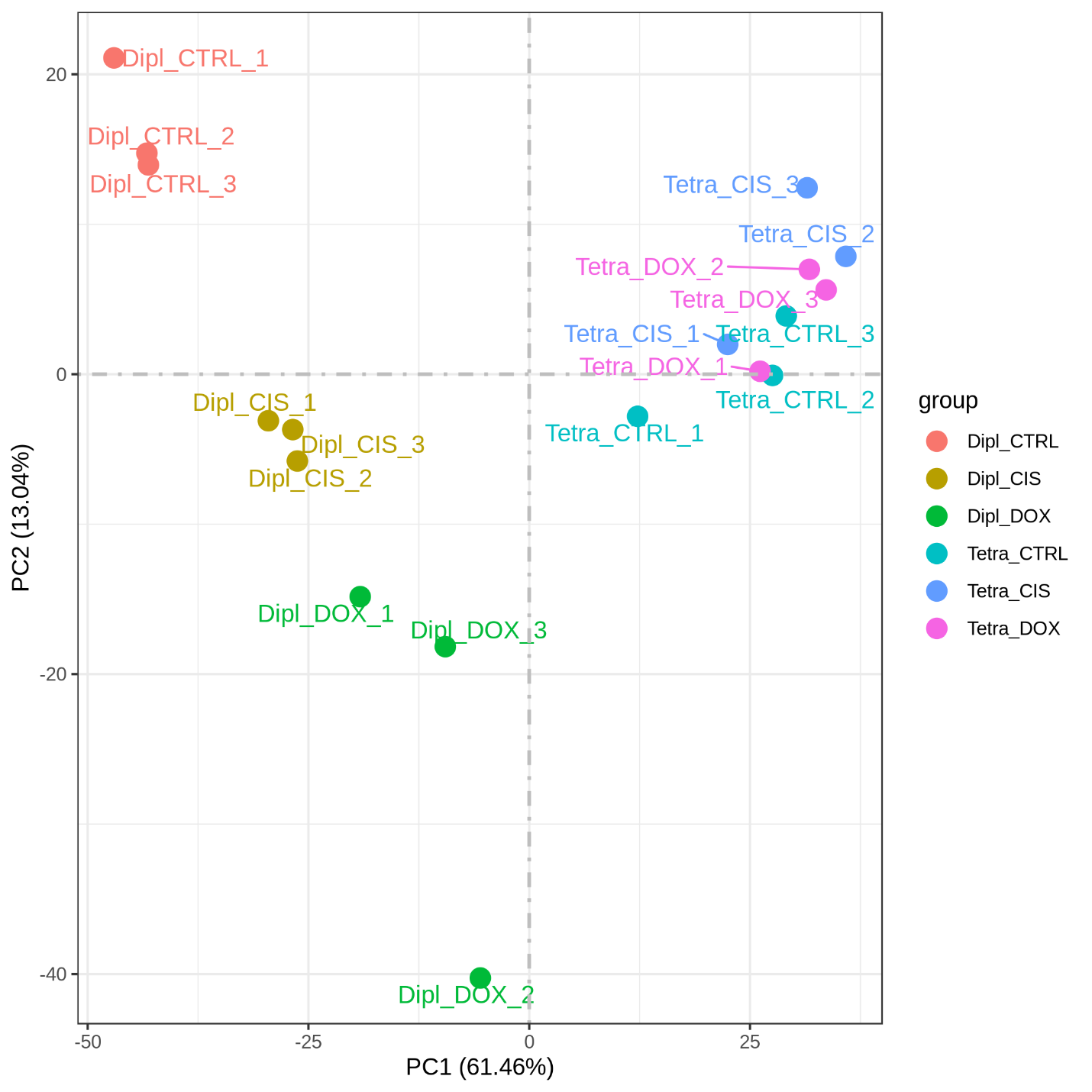

1. Department of Biochemistry and Molecular Biology; College of Medicine; Medical University of South Carolina, Charleston, South Carolina, 29425; United States of America
2. Department of Microbiology and Immunology; College of Medicine; Medical University of South Carolina, Charleston, South Carolina, 29425; United States of America
3. Department of Biology; College of Arts and Sciences; University of North Carolina, Chapel Hill, North Carolina, 27599; United States of America

   ^#^Equal contribution
